# Towards a machine-learning assisted diagnosis of psychiatric disorders and their operationalization in preclinical research: evidence from studies on addiction-like behaviour in individual rats

**DOI:** 10.1101/2021.10.11.463926

**Authors:** Kshitij S Jadhav, Benjamin Boury Jamot, Veronique Deroche-Gamonet, David Belin, Benjamin Boutrel

## Abstract

**Background:** Over the last few decades there is a progressive transition from a categorical to a dimensional approach to psychiatric disorders. Especially in case of substance use disorders, the increased interest in the individual vulnerability to transition from controlled to compulsive drug seeking and taking warrants the development of novel dimension-based objective diagnostic or stratification tools. Here we drew on a multidimensional preclinical model of addiction, namely the 3-criteria model, previously developed to identify the neurobehavioural basis of the individual vulnerability to switch from control to compulsive drug taking, to test the potential interest of a machine-learning assisted classifier objectively to identify individual subjects as vulnerable or resistant to addiction.

**Methods:** Large behavioural datasets from several of our previous studies on addiction-like behaviour for cocaine or alcohol were fed to a variety of machine-learning algorithms (each consisting of an unsupervised-clustering method combined with a supervised-prediction algorithm) to develop a classifier that identifies resilient and vulnerable rats with high precision and reproducibility irrespective of the cohort to which they belong.

**Results:** A classifier based on K-median or K-mean-clustering (for cocaine or alcohol, respectively) followed by Artificial Neural Networks emerged as a highly reliable and accurate tool to predict if a single rat is vulnerable or resilient to addiction. Thus, each of the rats previously characterized as displaying 0 criterion (i.e., resilient) or 3 criteria (i.e., vulnerable) in individual cohorts were correctly labelled by this classifier.

**Conclusion:** The present machine-learning-based classifier objectively labels single individuals as resilient or vulnerable to develop addiction-like behaviour in multisymptomatic preclinical models of cocaine or alcohol addiction-like behaviour in rats. This novel dimension-based classifier thereby increases the heuristic value and generalizability of these preclinical models while providing proof of principle for the deployment of similar tools for the future of diagnosis of psychiatric disorders.

## Introduction

### Background

There is a progressive transition from a categorical to a dimensional approach to psychiatric disorders (1–4). Meanwhile, a growing interest in the preclinical field of substance use disorder (SUD) is emerging, acknowledging the biobehavioural basis of the individual tendency to display aberrant, persistent, or compulsive, drug seeking and/or taking (5–10), warrants the development of new dimension-based objective diagnostic or stratification tools aiming at unveiling dimension-based, trans-nosological endophenotypes of vulnerability to SUDs.

Indeed, like for many psychiatric disorders, the presence of a triggering factor, such as exposure to a drug in the case of SUD, is not sufficient for the development of the several behaviours, which by the extreme nature of their manifestation alongside the continuum of their respective dimension, are characteristic of the pathology. This individual vulnerability to transition from controlled, recreational drug use to the compulsive drug seeking and taking that characterizes SUD (**Figure 1**) has long been suggested to stem from the interaction between environmental, psychological, neurobiological and behavioural factors (11–18). However, it is difficult to identify and study the biobehavioural basis of the factors conferring this vulnerability in humans, not least because such endeavours require the study of large populations across the lifetime in controlled conditions, with little if any opportunity to carry out the invasive manipulations that are necessary causally to identify the underlying neural and cellular mechanisms.

**Figure 1:**
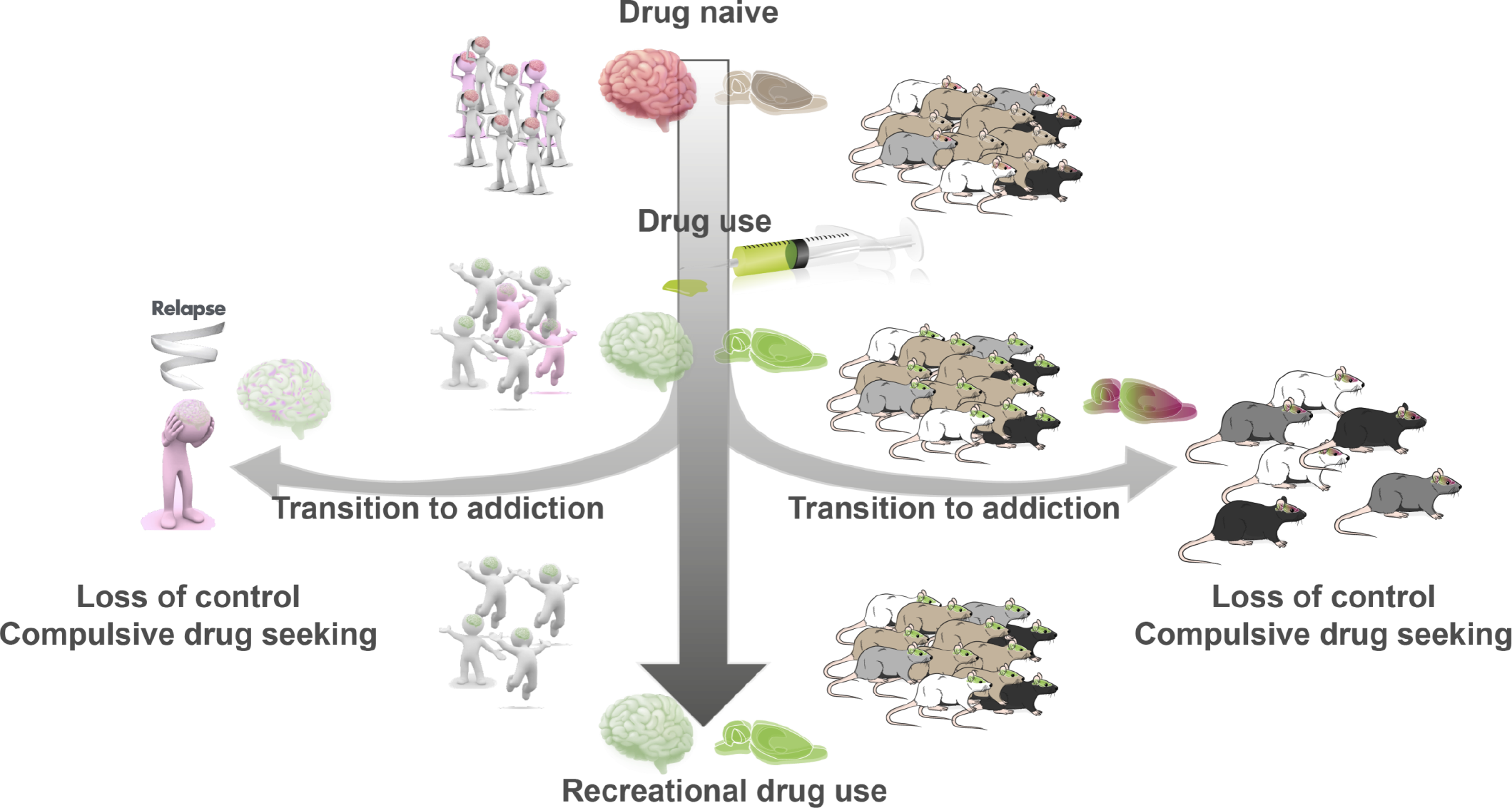
Individual trajectories in the development of substance use disorder: A translational perspective. Individuals who consume drugs, inadvertently take a risk since 10-40% of them eventually develop substance use disorder. While the majority maintain control over drug use, only a small proportion of individuals (humans/rats) progress to develop compulsive drug seeking which characterizes addiction. Recreational users will display drug-induced neurobiological changes (displayed here in green), but these changes, identified when recreational users are compared with drug naïve controls for instance, are not reflective of the adaptations to drug exposure that take place in the brain of vulnerable individuals (displayed here in pink), and which eventually mediate the transition to SUD and the associated high tendency to relapse. A frontier in addiction research is to understand what these addiction-specific adaptations are and what makes the brain of a vulnerable individual vulnerable. Over the past decades, it has been shown that similar differences exist in the tendency to develop addiction-like behaviour in rodents exposed to drug self-administration, and associated mechanisms identified in prospective longitudinal studies in rodents have systematically been verified in humans.

Over the past two decades, preclinical models have progressively evolved to incorporate the importance of these individual differences, thereby offering unique opportunities to overcome these limitations by using prospective longitudinal studies to investigate the psychological and neural basis of the vulnerability to develop addiction-like behaviour (19). Indeed, as in humans, all individual rats who regularly self-administer or seek addictive drugs do not necessarily lose control over drug intake and develop the persistent, compulsive drug-seeking and taking that characterizes substance use disorder (SUD) **(Figure-1)** (20). In this context, in the early 2000’s a multidimensional model of addiction was developed (21) based on the intersectionality of specific behavioural characteristics that are the operationalization of DSM-IV criteria (22), namely, increased motivation to take the drug, inability to refrain from drug-seeking and maintained drug use despite aversive consequences. This approach enables the identification of divergent trajectories of the transition from controlled to compulsive drug intake (**Figure-1**), in that only ~20% of a given population of outbred rats exposed to cocaine will eventually display the three behavioural criteria following a prolonged (>60 daily sessions) history of self-administration. Importantly, rats identified as displaying the 3-Criteria for addictive-behaviours also show an increased tendency to escalate their drug intake when access is illimited, and they are prone to relapse following abstinence (23), thereby displaying additional behavioural manifestations reminiscent of diagnostic criteria for which they were not selected. The construct and predictive validity of the 3criteria model was further substantiated as these differences between vulnerable and resilient rats are not due to a differential cocaine exposure since all rats self-administer the same amount of drug before being identified as 3 vs 0 criteria (21). However, while they do not take more cocaine, 3crit rats develop a binge-like pattern of intake that precedes the transition to addiction (23).

The 3-Criteria model has led to several breakthroughs in our understanding of the vulnerability to addiction which together represent a unique success story of translational research. This model first helped establish that impulsivity (24) and boredom susceptibility (25) confer a vulnerability to switch from controlled to compulsive cocaine intake. In contrast, both sensation seeking, assessed as a greater locomotor response to novelty, and sign tracking, which predict an increased tendency to initiate drug self-administration and to respond to drug-paired cues, respectively, were revealed to confer resilience to addiction-like behaviours (24, 26). These observations in rats have paved the way for studies in humans confirming that the factors associated with recreational cocaine use are dissociable from those specifically associated with the transition to SUD (27). The evidence of a causal relationship between a high impulsivity trait and the subsequent vulnerability to develop compulsive behaviours (24, 28) has far-reaching implications for our understanding of the neural basis of addiction (26, 29, 30) which the model helped reveal to be very different to the biological responses to drug exposure. Thus, the tendency to persist in drug taking despite adverse consequences is associated with a rigidity, not an exacerbation, as it is the case following single or repeated administrations of drugs, of drug-induced synaptic plasticity (31–33).

This multidimensional approach has since been applied to the study of the neural and psychological basis of the vulnerability to develop alcohol use disorder (AUD) (34, 35) or uncontrolled food seeking (36) illustrating the high translational value of novel preclinical models that encapsulate the multidimensional nature of SUD and the importance of focusing on the individual.

However, these procedures are all dependent on defining a threshold above which a behaviour is deemed maladaptive, but where the cursor should be placed on a continuum to consider a behaviour abnormal is a very challenging question, especially at a time of a transition from categorical to dimensional approaches. For example, in the 3-Criteria model, the threshold used for the scoring of each addiction-like criterion is determined by the physical properties of the distribution of the population for one of the three criteria, namely resistance to punishment (21, 24, 25, 37). The bimodal distribution of this dimension offers an objective threshold selection for the associated criterion, but its application to the two other criteria, inability to relinquish drug seeking even in the absence of the drug and high motivation for the drug, which both follow a log-normal distribution, relies on the assumption that a similar rupture in the continuum exists in them too, which is an inherent limitation. In addition, a distribution-based threshold selection to ascribe diagnostic scores puts too large an emphasis on the population to which each individual belongs, the physical properties of which eventually contributing almost as much as the individual characteristics themselves to its characterization as displaying addiction-like behaviour or resilience to addiction. This thereby precludes the determination of the vulnerability status of a given individual considered independently of a particular cohort, a limitation of the underlying approach which together with the associated need to train large cohorts at once for long periods of time may have hindered the widespread adoption of the 3-criteria model.

Recent developments in machine learning may offer unprecedented means to overcome the aforementioned limitations as they have been suggested to refine the classification of individuals, including psychiatric patients, within sub-groups with shared underlying endophenotypes, to tailor treatment strategies (38). These approaches also have an advantage over classical statistics (e.g. null hypothesis testing, ANOVAs), since they uncover substructures/subgroups in data without necessarily receiving specific instructions. Further, the models built using machine learning tools can also extrapolate patterns learnt from the data with which they are trained to entirely new data sets with individual precision.

Hence, here we used the 3-Criteria multidimensional models for cocaine or alcohol addiction, to test the potential of machine-learning assisted classifiers to identify individuals with or without addiction-like behaviour in drawing on diagnostic-relevant dimensions of addiction (20, 22). For this we subjected the individual scores in each of the three addiction-like behaviors to different clustering algorithms and then validated the labels using supervised prediction algorithms.

## Materials and Methods

### Data

For addiction-like behaviour for cocaine, data from 3 published papers (23–25) were pooled, representing a cohort of 88 individuals **(All_data_cocaine.csv)**. For addiction-like behaviour for alcohol, the data were pooled from 2 published (34, 35) and 1 unpublished experiment with a cohort of 150 rats **(All_data_alcohol.csv)**.

All analyses were processed using Python 3.8 using Numpy, Pandas and Scikit-learn packages as well as TensorFlow-Keras for deep learning methods (39, 40).

The active lever presses performed in each of the three behaviours (termed as raw data), namely, increased motivation to take the drug, inability to refrain from drug seeking and maintained drug use despite aversive consequences (compulsivity), were used as the three dimensions, injected in the algorithms.

The large datasets were split 50 times in 50 different training (67%) and test sets (33%) to avoid a cohort-driven bias in the clustering of individual rats **(Figure-2①)**.

**Figure 2:**
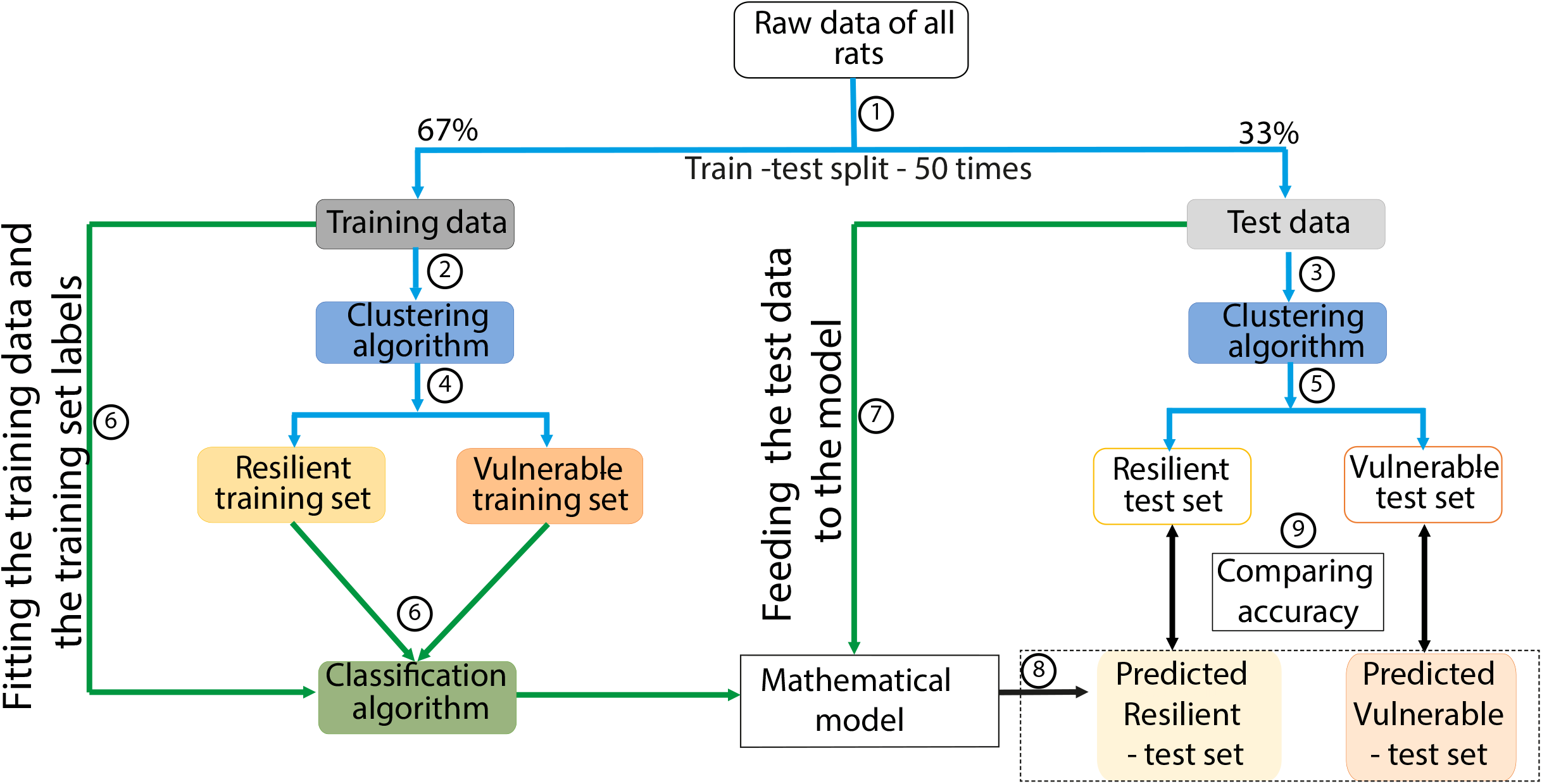
Workflow of the Machine Learning classifier. The steps are illustrated as numbers in the circles. Clustering algorithms used are Gaussian Mixture Method and K-mean/ K-median Clustering. Classification algorithms used are K nearest neighbor, Logistic Regression, Support Vector Machines and Artificial Neural Networks. The blue arrows indicate the clustering algorithms and the green arrows indicate the classification algorithms

### Algorithm

Individuals consuming drugs can be categorized as resilient or vulnerable to the development of SUD, the latter further being distributing along a clinical continuum of severity (18, 41, 42), suggesting that any population could be segregated in two clusters. Nevertheless, the optimal cluster number to be used in the classifier was determined by subjecting the 50 training and 50 test sets (i.e. 100 sets) to the Silhouette algorithm to inform the expected cluster number based on the actual experimental dataset **(cluster_number_cocaine.py, cluster_number_alcohol.py)** and the one most commonly informed by the Silhouette algorithm across 100 iterations, was included as an input in the clustering algorithms of the classifiers tested in the study.

Behavioural data of a single pair of TRAINING and TEST set was subjected to Unsupervised-Clustering algorithms **(Figure-2②,③)** (namely Gaussian Mixture Model (GMM) (43) or K-mean/K-median clustering (44)) **(SOM)** to determine resilient and vulnerable rats in both sets **(Figure-2④,⑤)**.

We used four Supervised Classification algorithms **(SOM)**, namely K nearest neighbour (KNN) (44), Logistic Regression (LR) (45), Support Vector Machines (SVM) (39) and Artificial Neural Networks (ANN) (46) (**Figure-3**) to fit the behavioural data of the TRAINING SET and the labels assigned by the clustering algorithm to generate a mathematical model that best explains the behavioural data and the labels of the rats in the TRAINING SET **(Figure-2⑥)**.

**Figure 3:**
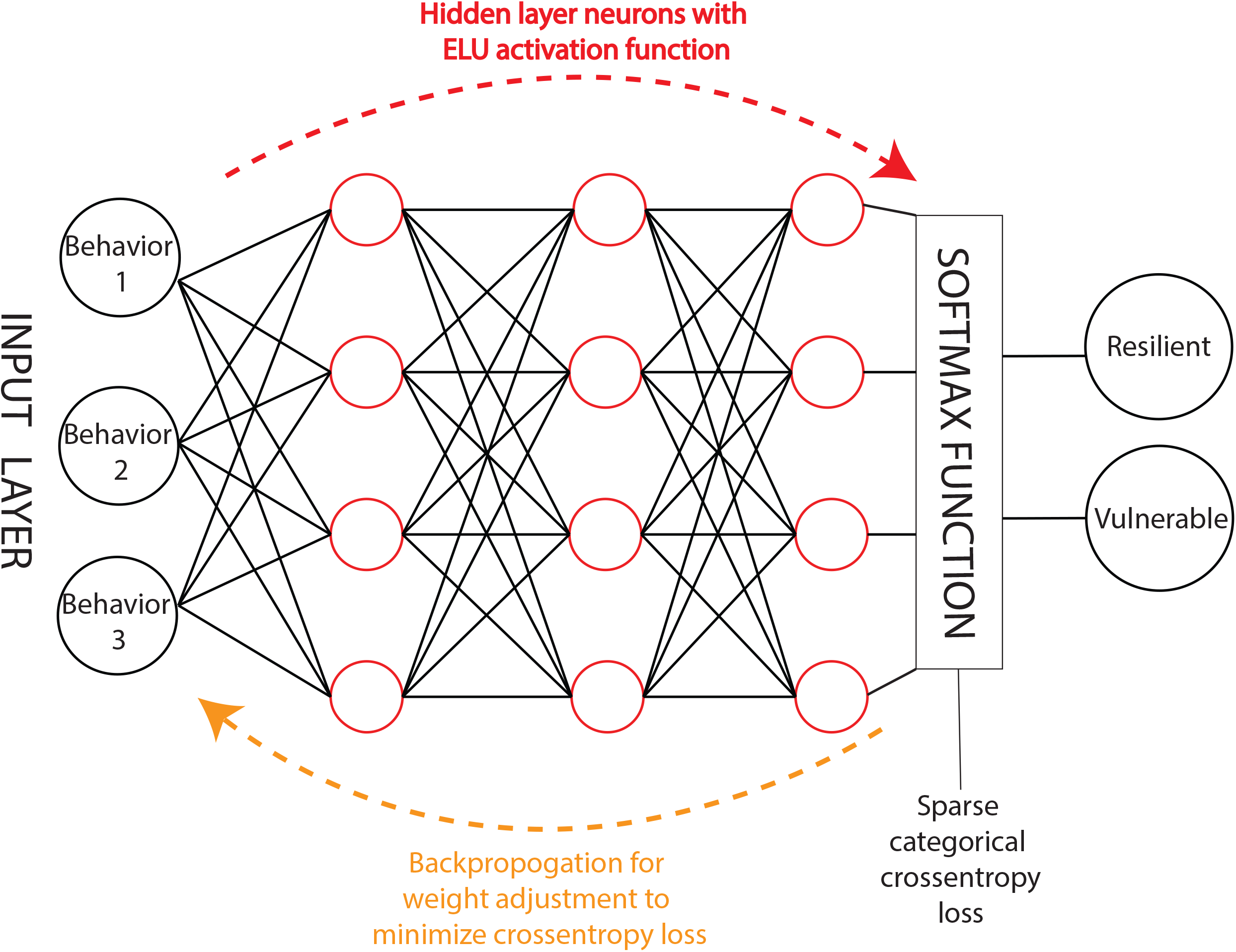
Illustration of an Artificial Neural Network. The hidden layer consists of neurons with the ELU (Exponential Linear Unit) activation function. Feed forward network entails multiple forward passes through the hidden layers. One forward pass consists of consecutive matrix multiplications at each layer by utilizing random weights to initialize the training which are then adjusted during backpropagation to minimize the cross entropy loss function.

For ANN, increasing numbers of hidden layers were used (5, 50 and 500), keeping the number of neurons in each layer constant, to test both a potential tendency to overfitting and the ability of the algorithm to accommodate larger sample sizes in the future. Then, to predict the labels of the rats belonging to the TEST set, their behavioural data were submitted to the mathematical model **(Figure-2⑦)** generated by each of the four Supervised Classification algorithms.

When submitted to these mathematical model, the behavioural data of the TEST SET are used to ascribe a resilient or vulnerable label to each rat of the TEST SET **(Figure-2⑧)** (39). Thus, each rat in the TEST set is ascribed two labels, one by the unsupervised-clustering algorithm and one by a particular supervised-prediction algorithm. The goal of this approach is to determine the Unsupervised Clustering–Supervised prediction combination that yields overlapping labels for the TEST SET rats **(Figure-2⑨)**.

The labels assigned to the TEST set rats by the clustering algorithm (considered here as true labels) and the predicted labels of the same rats by a supervised-prediction algorithm can be represented in a classification matrix (**Table-1**).

**Table 1:** Classification Matrix.

|  | Vulnerable (Supervised classification-based predictions- Test dataset) | Resilient (Supervised classification-based predictions- Test dataset) |
| --- | --- | --- |
| **Vulnerable (Unsupervised Clustering: test dataset)** | True Vulnerable (TV) | False Resilient (FR) |
| **Resilient (Unsupervised Clustering: test dataset)** | False Vulnerable (FV) | True resilient (TR) |

Accuracy ([TV+TR]/[TV+TR+FV+FR]), precision(TV/[TV+FV]), recall (TV/[TV+FR]) and ROC AUC (47) scores were calculated from the classification matrix (**SOM**).

Each pair of TRAINING and TEST SET were subjected to this pipeline four times, i.e. GMM-clustering followed by the four supervised-prediction algorithms. As mentioned previously, there were 50 pairs of TRAINING and TEST sets so that for each combination of GMM clustering-supervised prediction algorithm, fifty iterations were processed, resulting in fifty Accuracy, Precision, Recall and AUC ROC scores. Similarly, the same procedure was followed for K-median/K-mean clustering followed by four supervised prediction algorithms. Results are depicted as kernel density estimates of the probability density function of these fifty iterations for all four performance evaluation metrics for each combination of unsupervised clustering-supervised prediction algorithm.

## Results

For addiction-like behaviour for cocaine, Silhouette score revealed the optimal number of clusters was ‘2’ in 88% (Training sets–K-median clustering), 76% (Test sets–K-median clustering), 74% (Training sets– GMM clustering) and 76% (Test sets-GMM clustering), while the second most commonly suggested cluster number ranged from 3 to 6. Similarly, for addiction-like behaviour for alcohol, the optimal number of clusters was ‘2’ in 74% (Training sets–K-mean clustering), 78% (Test sets–K-mean clustering), 96% (Training sets–GMM clustering), 70% (Test sets–GMM clustering), while the second most commonly suggested cluster number ranged from 3-6. This analysis confirmed that 2 clusters should be used in subsequent analyses.

For addiction-like behaviour for cocaine, Kmedian-KNN, Kmedian-LR and Kmedian-SVM (**Kmedian_cocaine.py**) classifiers yielded similar scores (**Table 2A, Figure 4**) that were overall superior to GMM-KNN, GMM-LR and GMM-SVM (**GMM_cocaine.py**) classifiers with regard to median accuracy, precision, recall and ROC-AUC scores as well as the proportion of iterations reaching the top 10 percentile, respectively (**Table 2, Figure 4**). The Kmedian-ANN classifier and GMM-ANN classifier gave similar median accuracy and ROC-AUC scores and resulted in a similar proportion of these scores being in the top 10 percentile (**Table 2, Figure 4**).

**Figure 4:**
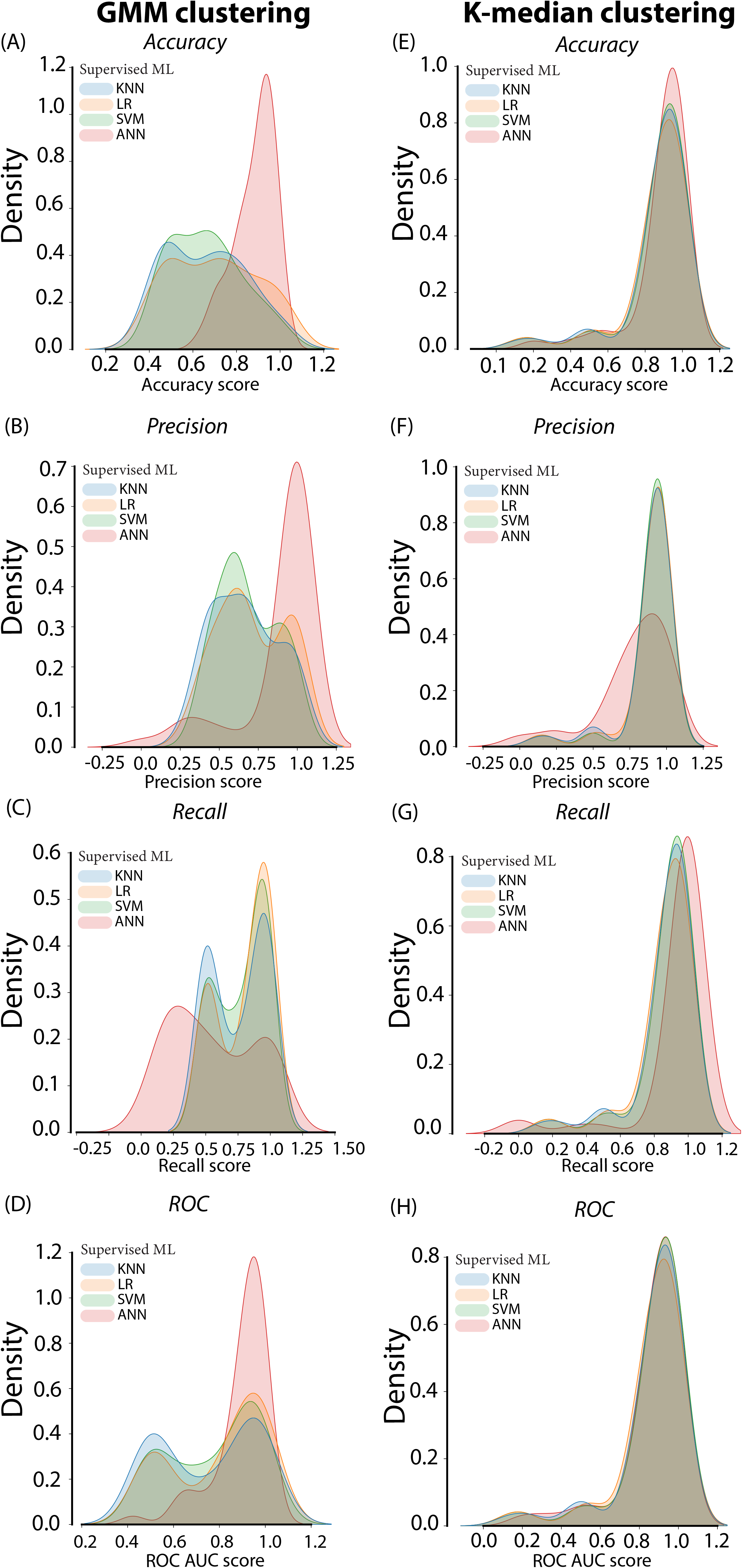
Performance evaluation metrics of the Machine Learning classifier of the Addiction-like behavior for cocaine in rats. Figure 4A-4D depict the Accuracy, Precision, Recall and ROC AUC score respectively of the GMM clustering based classifier followed by four Supervised Machine Learning algorithms. Figure 4E-4H depict the Accuracy, Precision, Recall and ROC AUC score respectively of the K-median clustering-based classifier followed by the four Supervised Machine Learning algorithms. GMM: Gaussian Mixture Method, ML: Machine Learning, KNN: K nearest neighbor, LR: Logistic Regression, SVM: Support Vector Machines, ANN: Artificial Neural Networks

**Table 2A:** Classifier based on K-median clustering followed by Supervised algorithm-based predictions for addiction-like behavior for cocaine.

|  | Accuracy |  | Precision |  | Recall |  | ROC AUC score |  |
| --- | --- | --- | --- | --- | --- | --- | --- | --- |
|  | Median | Scores in top 10 percentile | Median | Scores in top 10 percentile | Median | Scores in top 10 percentile | Median | Scores in top 10 percentile |
| KNN | 0.93 | 58% | 0.93 | 72% | 0.92 | 58% | 0.92 | 58% |
| LR | 0.92 | 56% | 0.93 | 72% | 0.91 | 52% | 0.91 | 52% |
| SVM | 0.93 | 60% | 0.92 | 76% | 0.93 | 60% | 0.93 | 60% |
| ANN<sup>5</sup> | 0.86 | 26% | 0.7 | 18% | 1 | 72% | 0.83 | 26% |
| ANN<sup>50</sup> | 0.93 | 52% | 0.80 | 36% | 1 | 88% | 0.9 | 48% |
| ANN<sup>500</sup> | 0.96 | 48% | 0.9 | 42% | 1 | 92% | 0.95 | 68% |

**Table 2B:** Classifier based on GMM clustering followed by Supervised algorithm-based predictions for addiction-like behavior for cocaine.

|  | Accuracy |  | Precision |  | Recall |  | ROC AUC score |  |
| --- | --- | --- | --- | --- | --- | --- | --- | --- |
|  | Median | Scores in top 10 percentile | Median | Scores in top 10 percentile | Median | Scores in top 10 percentile | Median | Scores in top 10 percentile |
| KNN | 0.67 | 16% | 0.75 | 28% | 0.88 | 48% | 0.88 | 48% |
| LR | 0.73 | 6% | 0.67 | 22% | 0.87 | 46% | 0.87 | 46% |
| SVM | 0.68 | 10% | 0.66 | 22% | 0.89 | 48% | 0.89 | 48% |
| ANN<sup>5</sup> | 0.9 | 34% | 1 | 52% | 0.36 | 22% | 0.84 | 42% |
| ANN<sup>50</sup> | 0.93 | 60% | 1 | 74% | 0.67 | 38% | 0.94 | 62% |
| ANN<sup>500</sup> | 0.93 | 56% | 1 | 80% | 0.6 | 24% | 0.96 | 74% |

GMM: Gaussian mixture model, SML: Supervised Machine Learning, KNN: K nearest neighbor, LR: Logistic Regression, SVM: Support Vector Machines, ANN: Artificial Neural Networks. ANN<sup>5</sup>, ANN<sup>50</sup>, ANN<sup>500</sup>: numbers indicate the number of hidden layers of the ANN

For addiction-like behaviour for alcohol, Kmean-KNN, Kmean-LR, Kmean-SVM and Kmean-ANN classifiers yielded similar scores for all the performance evaluation metrics (**Table 3A, Figure 5**) that were overall superior to those obtained by GMM-KNN, GMM-LR, GMM-SVM and GMM-ANN classifiers (**Table 3, Figure 5**).

**Figure 5:**
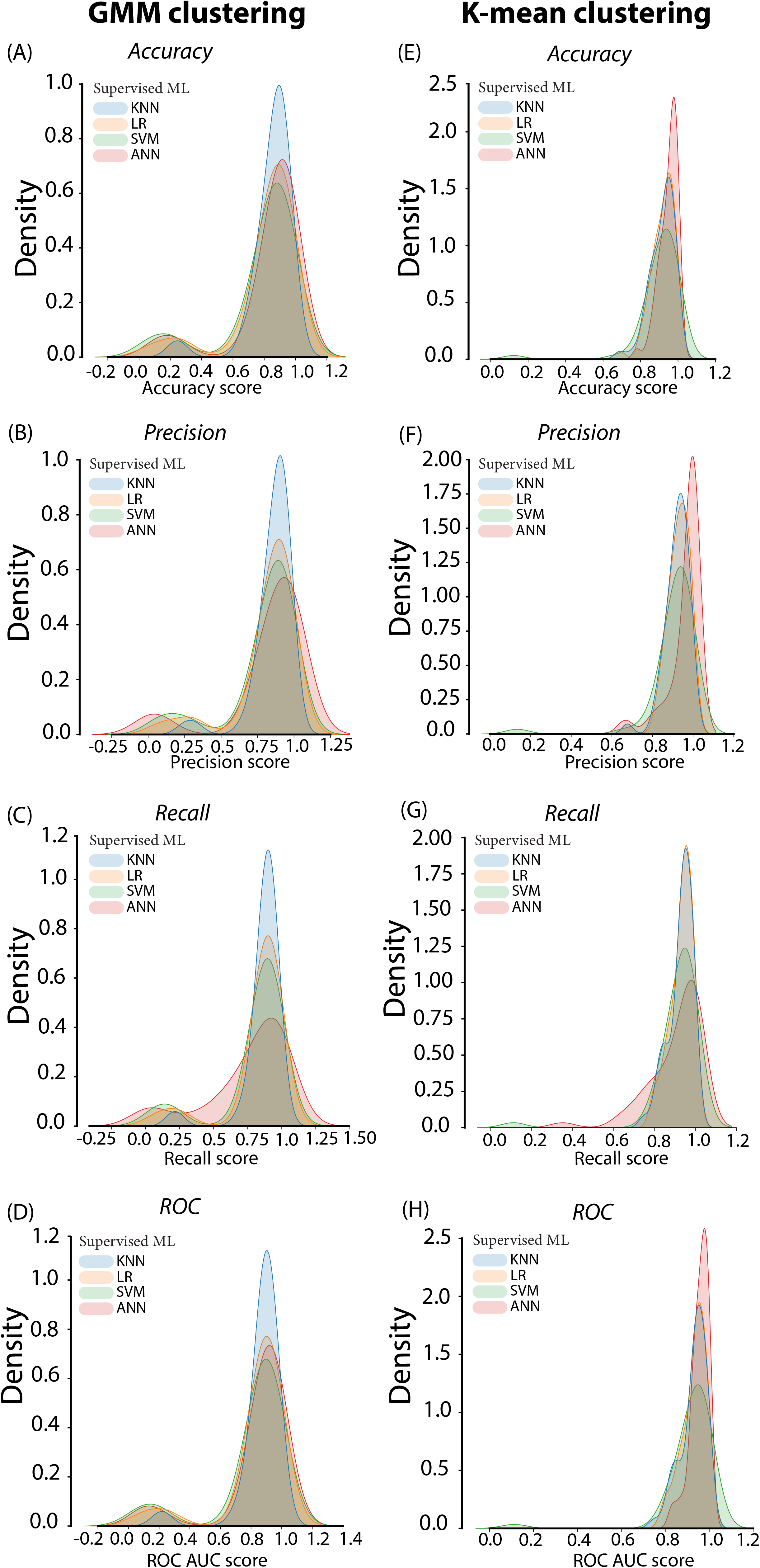
Performance evaluation metrics of the Machine Learning classifier of the Addiction-like behavior for alcohol in rats. Figure 5A-5D depict the Accuracy, Precision, Recall and ROC AUC score respectively of the GMM clustering based classifier followed by four Supervised Machine Learning algorithms. Figure 5E-5H depict the Accuracy, Precision, Recall and ROC AUC score respectively of the K-mean clustering-based classifier followed by four Supervised Machine Learning algorithms. GMM: Gaussian Mixture Method, ML: Machine Learning, KNN: K nearest neighbor, LR: Logistic Regression, SVM: Support Vector Machines, ANN: Artificial Neural Networks

**Table 3A:** Classifier based on K-mean clustering followed by Supervised algorithm-based predictions for addiction-like behavior for alcohol.

|  | Accuracy |  | Precision |  | Recall |  | ROC AUC score |  |
| --- | --- | --- | --- | --- | --- | --- | --- | --- |
|  | Median | Scores in top 10 percentile | Median | Scores in top 10 percentile | Median | Scores in top 10 percentile | Median | Scores in top 10 percentile |
| KNN | 0.92 | 66% | 0.93 | 68% | 0.94 | 78% | 0.94 | 78% |
| LR | 0.93 | 62% | 0.93 | 68% | 0.94 | 72% | 0.94 | 72% |
| SVM | 0.93 | 62% | 0.93 | 68% | 0.93 | 76% | 0.93 | 76% |
| ANN<sup>5</sup> | 0.92 | 60% | 0.93 | 60% | 0.94 | 60% | 0.93 | 66% |
| ANN<sup>50</sup> | 0.96 | 76% | 1 | 74% | 0.94 | 60% | 0.96 | 88% |
| ANN<sup>500</sup> | 0.96 | 82% | 1 | 86% | 0.97 | 64% | 0.97 | 90% |

**Table 3B:** Classifier based on GMM clustering followed by Supervised algorithmbased predictions for addiction-like behavior for alcohol.

|  | Accuracy |  | Precision |  | Recall |  | ROC AUC score |  |
| --- | --- | --- | --- | --- | --- | --- | --- | --- |
|  | Median | Scores in top 10 percentile | Median | Scores in top 10 percentile | Median | Scores in top 10 percentile | Median | Scores in top 10 percentile |
| KNN | 0.87 | 40% | 0.90 | 50% | 0.89 | 48% | 0.89 | 48% |
| LR | 0.87 | 42% | 0.89 | 46% | 0.89 | 46% | 0.89 | 46% |
| SVM | 0.86 | 40% | 0.88 | 46% | 0.89 | 44% | 0.89 | 44% |
| ANN<sup>5</sup> | 0.87 | 40% | 0.88 | 46% | 0.85 | 32% | 0.89 | 46% |
| ANN<sup>50</sup> | 0.88 | 36% | 0.91 | 56% | 0.82 | 38% | 0.89 | 44% |
| ANN<sup>500</sup> | 0.92 | 54% | 0.91 | 64% | 0.9 | 50% | 0.92 | 60% |

GMM: Gaussian mixture model, SML: Supervised Machine Learning, KNN: K nearest neighbor, LR: Logistic Regression, SVM: Support Vector Machines, ANN: Artificial Neural Networks. ANN<sup>5</sup>, ANN<sup>50</sup>, ANN<sup>500</sup>: numbers indicate the number of hidden layers of the ANN

The performance of the Kmedian- and GMM-ANN or Kmean- and GMM-ANN classifiers was further improved by an increase in the number of hidden layer neurons used in the ANN (**Table 2 and 3**). The ensuing increase in accuracy thereby demonstrates the ability of the Kmedian/Kmean-ANN classifiers to accommodate larger sample sizes and/or more dimensions, a feature that is not reflective simply of overfitting because of the back propagation and early stopping processes included in the ANN (48).

Together these results demonstrate that a classifier based on K-median/K-mean followed by ANN is the most robust and future- and dimension expansion-proof approach accurately to predict whether a single rat is vulnerable or resilient as assessed in our multisymptomatic model with great heuristic value with regards to the clinical definition of SUD **(Figure-6)**.

**Figure 6:**
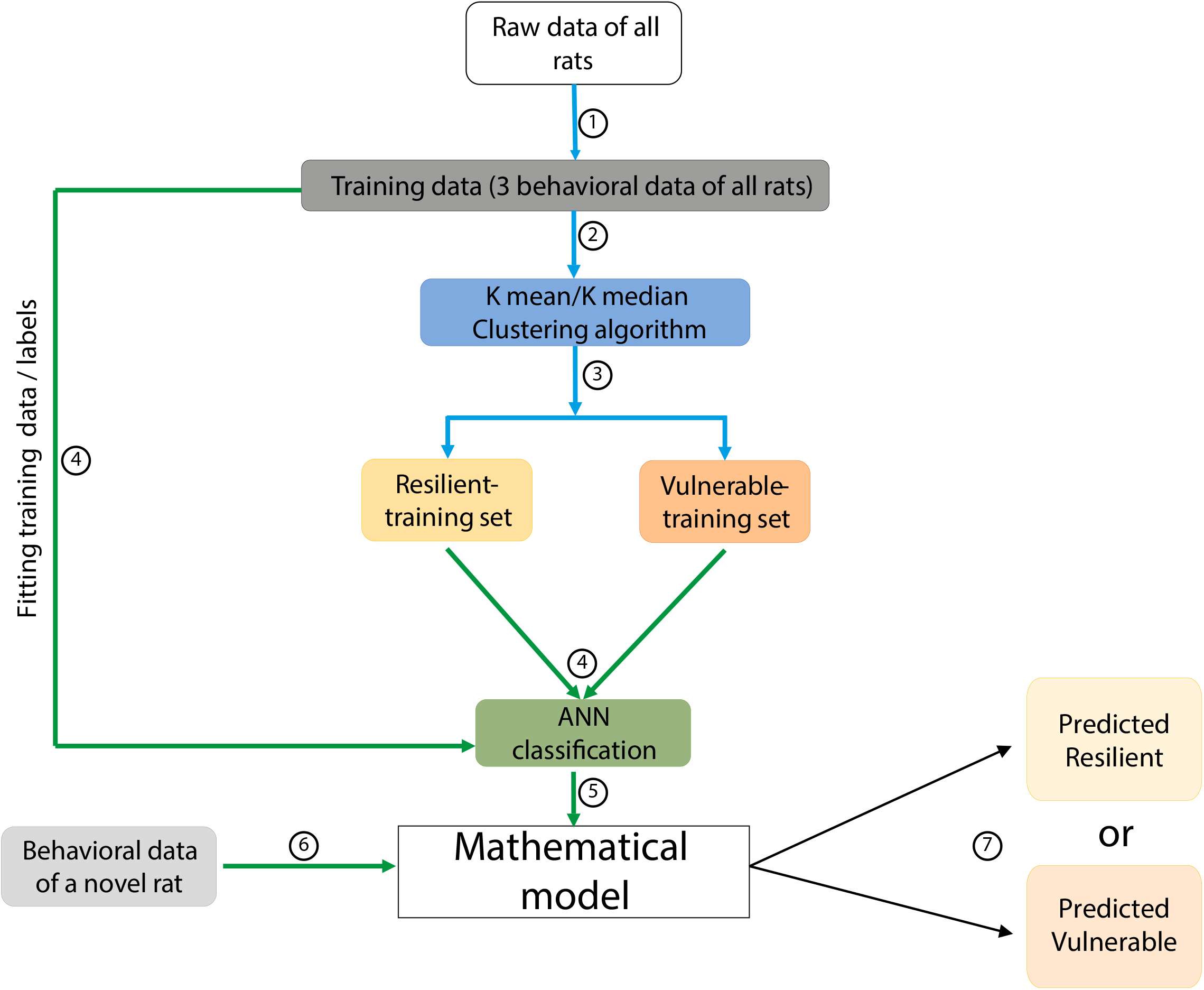
Flowchart of the classification of any future rat as resilient or vulnerable to SUD. The steps are illustrated as numbers in the circles. Having established that the classifier based on K-median/K-mean clustering followed by ANN gives the best predictive accuracy, the addiction vulnerability status of a single rat irrespective of the cohort it is trained with. The blue arrows indicate the clustering algorithms and the green arrows indicate the classification algorithms ANN: Artificial Neural Network

To cross-validate the classifier, the entire datasets related to cocaine and alcohol addiction-like behaviour (n=88, n=150, respectively) were subjected to the K-median (**All_cocaine_Kmedian.py**)/K-mean clustering (**All_alcohol_Kmean.py**). All the rats originally characterized as 0 or 3crit in their respective cohort were correctly labelled as Resilient or Vulnerable, respectively, revealing an absolute intersection (**Table-4**) (**Cocaine_crit_correspondence.xlsx,Alcohol_crit_correspondence.xlsx**).

**Table 4A:** Addiction-like behavior for cocaine.

|  | 0Crit in the original cohort | 3Crit in the original cohort |
| --- | --- | --- |
| Resilient (K median clustering) | 36 | 0 |
| Vulnerable (K median clustering) | 0 | 15 |

**Table 4B:** Addiction-like behavior for alcohol.

|  | 0Crit in the original cohort | 3Crit in the original cohort |
| --- | --- | --- |
| Resilient (K mean clustering) | 64 | 0 |
| Vulnerable (K mean clustering) | 0 | 18 |

Finally, in order to establish the predictive potential of the classifiers we developed, we applied them to a completely new dataset (26) that consists of the 3criteria behavioural scores of a cohort of 36 rats housed either in a standard (2 individuals in a standard cage), or an enriched environment. While replicating previous findings that an environmental enrichment decreases the tendency to self-administer cocaine (49, 50), this study demonstrated that rats housed in an enriched environment were more vulnerable to develop addiction-like behaviour than rats raised in a standard environment (26) in that all the 3crit rats identified in this heterogeneous cohort came from the former. In line with the original study, none of the rats from the standard housing group were labelled as vulnerable by the classifiers, while those identified as vulnerable overlapped with 100% accuracy with those identified as 3crit that came from the enriched environment (**EEES.xlsx**) (26).

## Discussion

The next frontier in addiction research lies in understanding the environmental, psychological and biological mechanisms that mediate, in vulnerable individuals, the transition from controlled drug intake to the compulsive seeking and taking characteristic of SUD. Behavioural procedures that enable the study, under controlled conditions, of individual trajectories from a drug naïve state to addiction-like behaviour development, over the course of drug self-administration (21, 23, 24, 26, 29, 34, 35) have only started to demonstrate their utility in our understanding of the mechanisms of the individual vulnerability to addiction. These procedures have hitherto been limited by a lack of objective diagnosis strategy that is not influenced by the physical data-distribution properties of the cohort to which a single individual belongs. This results in the unwarranted need to train large cohorts of animals at any given time and detracts the approach from the individual-centred diagnosis in humans.

In this study, we drew on large datasets produced over the past decade to develop new machine-learning assisted classifiers for cocaine or alcohol addiction-like behaviour that characterize with high accuracy single individuals, irrespective of the cohort to which they belong, as resilient or vulnerable.

The role of clustering algorithms is to identify data-points in a multidimensional space that are closer to one another than they are to any other data point in the cloud (51). In many real-life situations, the labels of such data-points are obvious, e.g., Males vs Females for biological differences, or voted for or against Brexit. In these situations, data clustering is not necessary. However, ascribing labels, such as those to determine if an individual meets the criteria of addiction-like behaviour, cannot be informed by a natural dichotomic population segregation. This requires to structure a multidimensional space in delineated sub-spaces which can be used to ascribe a specific label to each individual constituent of the cluster and to train supervised classification algorithms in order to successfully predict the label, i.e., the specific cluster to which they most likely belong of a single individual whose data have never been used to train the classification algorithm.

The first step of such an algorithm development was to objectively determine the cluster number to structure the multidimensional cloud to accommodate the physical properties of the data and the objective of the classifier. In real life, individuals can be categorized as vulnerable or resilient, thereby suggesting that any experimental population could be segregated into two clusters (18, 41, 42). Nevertheless, a data-driven approach was used to ensure that such dichotomy was present in the experimental datasets. The Silhouette algorithm ran on all the datasets used here systematically revealed that the multidimensional space of the datasets was predominantly structured around two clusters, an outcome that is compatible with the prerequisite of the algorithm: to segregate two subpopulations from heterogeneous groups, namely vulnerable and resilient individuals. This also provided an unbiased threshold for the various cluster analyses (GMM and K-median/K-mean) used in the several potential classifiers tested in this study.

Identification of resilient or vulnerable rats in the 3-Criteria model was hitherto based on the bimodal distribution of each population for resistance to punishment (21, 24, 34) which comprises a large log-normally distributed subpopulation of non-compulsive rats (60-70% population) tailed by an independent, normally distributed population of compulsive rats (30-40% population). Since GMM-clustering can fit bi/multimodal data distributions (52), it was originally used to assimilate such physical property on which depends the selection threshold for addiction-like behaviour. However, the GMM-based classifier did not yield outputs superior to the K-median/K-mean based classifier. This surprising outcome can be due to the fact that a GMM classifier, in contrast with the strategy we developed to apply the 30-40% threshold to the other two criteria, each characterized by a log-normal distribution, uses differential densities across quartiles in each variable independently to develop the classifier.

K-median/K-mean clustering, which is based on the Euclidean distance between the data-points in a three-dimensional vector space that plots the number of responses along three axes representing three different psychological constructs, was revealed to be the superior clustering method to accurately and consistently ascribe labels of resilience vs vulnerability.

Not only are K-median/K-mean algorithms easy to implement, but they are scalable and can be used to separate non-linearly separable data. These properties were exploited to develop a robust and universal classifier. Thus, the same clustering algorithms were applied to fifty independent sets drawn from a large dataset comprising, in the case of cocaine addiction-like behaviour, data from experiments carried out in different laboratories, on different strains (Sprague-Dawley (23, 25) or Lister-Hooded (24) or Wistar rats (34, 35)) that differ in addiction relevant traits (53) and using different instrumental responses (nose-pokes (23, 25) or lever presses (24)). The ability of the classifier to survive randomization tests and to generalize across response modalities and strains demonstrates its potential use across a large repertoire of experimental idiosyncrasies that may reflect the behavioural heterogeneities observed by clinicians when a diagnosis is warranted. Further, the ability of the Kmedian/Kmean-supervised algorithm classifiers to accurately identify rats as being vulnerable to addiction or resilient from a completely different dataset generated with a heterogeneous cohort exposed to very different housing conditions to those used in the experiments used for the training datasets indicates that these tools can be deployed across many experimental conditions.

Nevertheless, the same classifier could not be generalized from addiction-like behaviour for one drug to another. While K-median clustering-based algorithm used for addiction-like behaviour for cocaine systematically yielded the right vulnerability/resilient labels, even when applied to a dataset never used in its development (the enriched environment experiment (26)), it was sub-optimal in the case of addiction-like behaviour for alcohol, the best classifier for which was based on K-mean clustering. The lack of generalizability of a given classifier across drugs is a further evidence of construct and predictive validity since AUD and SUD are independent diagnoses in humans and they have long been shown to involve different psychological and neurobiological mechanisms (54). In addition, while the 3-Criteria model for cocaine relies on the assessment of compulsive cocaine intake (consummatory conflated with preparatory responses as it is the case under fixed-ratio schedules of reinforcement) (19), the 3-Criteria model for alcohol is based on the assessment of the compulsive nature of a seeking response in a chained schedule where lever pressing results in the procurement of alcohol, the ensuing consumption of which occurs in a dedicated magazine, involving a set of behavioural responses independent of the instrumental component of the chain. Considering how neurally and psychologically dissociable preparatory and consummatory responses are (55, 56), it was not expected that a single classifier could be used across measures of compulsive taking and seeking. However, it will be interesting to test in future studies if the alcohol-specific K-mean classifier can be applied to compulsive cocaine seeking data, as measured in seeking-taking heterogeneous chained schedules of reinforcement, which dissociated physically and spatially, seeking response from taking/consummatory responses for intravenously administered drug (57, 58).

Irrespective of the outcomes of these future studies, a one-fit-all approach is not an optimal expectation. One important avenue for future research is to identify mathematical tools that will enable the introduction of dimensionality within the categories that are now identified accurately with the K-mean/K-median classifiers. The 3-Criteria model was designed to have construct validity with regards to the diagnosis strategy of DSM-IV (22), i.e., prior to the development of the RDoC (1). Nevertheless, the approach we had then developed embedded a dimensional aspect, in that rats were not only stratified as showing 0 criterion or 3 criteria (deemed resilient and showing addiction-like behaviour, respectively), but 30-40% of any population was also stratified as showing 1 or 2 criteria, with 1crit and 2crit rats being considered similar to 0crit and 3crit respectively (34, 59, 60).

Since all the resilient rats identified by our classifier included 0crit, most of the 1crit and no 3crit rats, whereas all the rats identified as vulnerable included 3crit and most of the 2crit but no 0crit rats (**Cocaine_crit_correspondence.xlsx,Alcohol_crit_correspondence.xlsx**), it can be suggested that the present classifier does not yet provide the dimensional granularity necessary to distinguish several levels of severity (2crit vs 3crit) within the vulnerable population warranting further research to determine whether the addition of endophenotypes (24, 25, 34, 35) to our classifiers will enable them to fully comply with the dimensional nature of the debilitating condition that is SUD. This could contribute to the several initiatives to identify clinically relevant subtypes of SUD (61) through cluster analysis of patients to better capture the clinical heterogeneity (62–66) with the aim to advance personalized medicine (67, 68).

## Conclusion

The present machine learning-based classifiers represent a unique tool to objectively identify whether a single experimental subject is resilient or vulnerable for cocaine or alcohol addiction-like behaviour (**Figure-6**). The ability conferred by such a tool to consider a single individual irrespective of the experimental cohort to which it belongs (and the associated experimental conditions) bridges a new frontier in the study of the individual vulnerability to develop SUD, bringing the focus back on the individual, as it is the case in humans. It can be boldly envisioned that, with the advent of large data sets in humans from imaging, genomics and proteomic approaches a successful back-translation strategy could see the application of such machine learning-assisted tools to a personalized diagnosis of clinical populations.

## Supporting information

Alcohol_crit_correspondence.xlsx

Cocaine_crit_correspondence.xlsx

EEES.xlsx

All_alcohol_Kmean.py

All_cocaine_Kmedian.py

cluster_numbers_alcohol.py

cluster_numbers_cocaine.py

GMM_alcohol.py

GMM_cocaine.py

Kmean_alcohol.py

Kmedian_cocaine.py

SOM

All_data_alcohol.csv

All_data_cocaine.csv

## Conflict of Interest

The authors declare no competing financial interests.

## Funding

DB is supported by a Programme Grant from the Medical Research Council to Barry Everitt, DB, Amy Milton, Jeffrey Dalley and Trevor Robbins (MR/N02530X/1). VDG was supported by ANR-13-NEUR-0002-01 (ERA-NET NEURON II), ANR-10-LABX-43 (LABEX BRAIN), ANR-10-EQX-008-1 (EquipEx OptoPath™), INSERM and Université de Bordeaux. BB is supported by a Swiss National Science Foundation grant (310030_185192)

KSJ is the recipient of the Swiss Government Excellence Fellowship (2015-2018), Doc mobility Fellowship (2018-2019) and Early Post Doc mobility Fellowship (2021-2022) from the Swiss National Science Foundation.

## Author contribution

KSJ developed the algorithms and wrote the codes. BBJ provided substantial technical inputs for the code. KSJ, DB, VDG and BB wrote the manuscript

## Acknowledgement for assistance

BB and KSJ would like to thank Dr Ivana Arsic, (Data Scientist, Center for Psychiatric Neuroscience, Department of Psychiatry, Lausanne University Hospital, Switzerland) for her inputs on the algorithm.

KSJ would like to thank Mr. Dhaval Joshi, PhD student in the Belin lab for critical insights that helped to refine the algorithms and the annotating the code in the .py files.

## Notes

### Competing Interest Statement

The authors have declared no competing interest.

## References

1. Brooks SJ, Lochner C, Shoptaw S, Stein DJ (2017): Using the research domain criteria (RDoC) to conceptualize impulsivity and compulsivity in relation to addiction. Prog Brain Res. 235:177–218.

2. Insel TR (2014): The NIMH Research Domain Criteria (RDoC) Project: precision medicine for psychiatry. Am J Psychiatry. 171:395–397.

3. Woody ML, Gibb BE (2015): Integrating NIMH Research Domain Criteria (RDoC) into Depression Research. Curr Opin Psychol. 4:6–12.

4. Ford JM, Morris SE, Hoffman RE, Sommer I, Waters F, McCarthy-Jones S, et al. (2014): Studying hallucinations within the NIMH RDoC framework. Schizophr Bull. 40 Suppl 4:S295–304.

5. Zhukovsky P, Puaud M, Jupp B, Sala-Bayo J, Alsio J, Xia J, et al. (2019): Withdrawal from escalated cocaine self-administration impairs reversal learning by disrupting the effects of negative feedback on reward exploitation: a behavioral and computational analysis. Neuropsychopharmacology. 44:2163–2173.

6. McNamara R, Dalley JW, Robbins TW, Everitt BJ, Belin D (2010): Trait-like impulsivity does not predict escalation of heroin self-administration in the rat. Psychopharmacology (Berl). 212:453–464.

7. Augier E, Barbier E, Dulman RS, Licheri V, Augier G, Domi E, et al. (2018): A molecular mechanism for choosing alcohol over an alternative reward. Science. 360:1321–1326.

8. Pohorala V, Enkel T, Bartsch D, Spanagel R, Bernardi RE (2021): Sign-and goal-tracking score does not correlate with addiction-like behavior following prolonged cocaine self-administration. Psychopharmacology (Berl).

9. Cannella N, Cosa-Linan A, Takahashi T, Weber-Fahr W, Spanagel R (2020): Cocaine addicted rats show reduced neural activity as revealed by manganese-enhanced MRI. Sci Rep. 10:19353.

10. Lenoir M, Serre F, Cantin L, Ahmed SH (2007): Intense sweetness surpasses cocaine reward. PLoS One. 2:e698.

11. Anthony J, Warner L, Kessler R (1994): Comparative epidemiology of dependence on tobacco, alcohol, controlled substances, and inhalants: Basic findings from the National Comorbidity Survey. Experimental and Clinical Psychopharmacology. 2:244–268.

12. Grant BF, Stinson FS, Harford TC (2001): Age at onset of alcohol use and DSM-IV alcohol abuse and dependence: a 12-year follow-up. J Subst Abuse. 13:493–504.

13. Grant BF, Stinson FS, Dawson DA, Chou SP, Dufour MC, Compton W, et al. (2004): Prevalence and co-occurrence of substance use disorders and independent mood and anxiety disorders: results from the National Epidemiologic Survey on Alcohol and Related Conditions. Arch Gen Psychiatry. 61:807–816.

14. Conway KP, Swendsen JD, Rounsaville BJ, Merikangas KR (2002): Personality, drug of choice, and comorbid psychopathology among substance abusers. Drug Alcohol Depend. 65:225–234.

15. Swendsen J, Conway KP, Degenhardt L, Dierker L, Glantz M, Jin R, et al. (2009): Socio-demographic risk factors for alcohol and drug dependence: the 10-year follow-up of the national comorbidity survey. Addiction. 104:1346–1355.

16. Swendsen J, Conway KP, Degenhardt L, Glantz M, Jin R, Merikangas KR, et al. (2010): Mental disorders as risk factors for substance use, abuse and dependence: results from the 10-year follow-up of the National Comorbidity Survey. Addiction. 105:1117–1128.

17. Ersche KD, Jones PS, Williams GB, Turton AJ, Robbins TW, Bullmore ET (2012): Abnormal brain structure implicated in stimulant drug addiction. Science. 335:601–604.

18. Ersche KD, Meng C, Ziauddeen H, Stochl J, Williams GB, Bullmore ET, et al. (2020): Brain networks underlying vulnerability and resilience to drug addiction. Proc Natl Acad Sci U S A. 117:15253–15261.

19. Belin-Rauscent A, Fouyssac M, Bonci A, Belin D (2016): How Preclinical Models Evolved to Resemble the Diagnostic Criteria of Drug Addiction. Biol Psychiatry. 79:39–46.

20. (2013): American Psychiatric Association: Diagnostic and Statistical Manual of Mental Disorders. 5th ed ed. Arlington: American Psychiatric Association.

21. Deroche-Gamonet V, Belin D, Piazza PV (2004): Evidence for addiction-like behavior in the rat. Science. 305:1014–1017.

22. (1994): Diagnostic and statistical manual of mental disorders: DSM-IV. Washington DC: American Psychiatric Association.

23. Belin D, Balado E, Piazza PV, Deroche-Gamonet V (2009): Pattern of intake and drug craving predict the development of cocaine addiction-like behavior in rats. Biol Psychiatry. 65:863–868.

24. Belin D, Mar AC, Dalley JW, Robbins TW, Everitt BJ (2008): High impulsivity predicts the switch to compulsive cocaine-taking. Science. 320:1352–1355.

25. Belin D, Berson N, Balado E, Piazza PV, Deroche-Gamonet V (2011): High-novelty-preference rats are predisposed to compulsive cocaine self-administration. Neuropsychopharmacology. 36:569–579.

26. Fouyssac M, Puaud M, Ducret E, Marti-Prats L, Vanhille N, Ansquer S, et al. (2021): Environment-dependent behavioral traits and experiential factors shape addiction vulnerability. Eur J Neurosci. 53:1794–1808.

27. Ersche KD, Turton AJ, Pradhan S, Bullmore ET, Robbins TW (2010): Drug addiction endophenotypes: impulsive versus sensation-seeking personality traits. Biol Psychiatry. 68:770–773.

28. Ansquer S, Belin-Rauscent A, Dugast E, Duran T, Benatru I, Mar AC, et al. (2014): Atomoxetine decreases vulnerability to develop compulsivity in high impulsive rats. Biol Psychiatry. 75:825–832.

29. Besson M, Pelloux Y, Dilleen R, Theobald DE, Lyon A, Belin-Rauscent A, et al. (2013): Cocaine modulation of frontostriatal expression of Zif268, D2, and 5-HT2c receptors in high and low impulsive rats. Neuropsychopharmacology. 38:1963–1973.

30. Dalley JW, Fryer TD, Brichard L, Robinson ES, Theobald DE, Laane K, et al. (2007): Nucleus accumbens D2/3 receptors predict trait impulsivity and cocaine reinforcement. Science. 315:1267–1270.

31. Pascoli V, Terrier J, Espallergues J, Valjent E, O’Connor EC, Luscher C (2014): Contrasting forms of cocaine-evoked plasticity control components of relapse. Nature. 509:459–464.

32. Ungless MA, Whistler JL, Malenka RC, Bonci A (2001): Single cocaine exposure in vivo induces long-term potentiation in dopamine neurons. Nature. 411:583–587.

33. Kasanetz F, Deroche-Gamonet V, Berson N, Balado E, Lafourcade M, Manzoni O, et al. (2010): Transition to addiction is associated with a persistent impairment in synaptic plasticity. Science. 328:1709–1712.

34. Jadhav KS, Magistretti PJ, Halfon O, Augsburger M, Boutrel B (2017): A preclinical model for identifying rats at risk of alcohol use disorder. Sci Rep. 7:9454.

35. Jadhav KS, Peterson VL, Halfon O, Ahern G, Fouhy F, Stanton C, et al. (2018): Gut microbiome correlates with altered striatal dopamine receptor expression in a model of compulsive alcohol seeking. Neuropharmacology. 141:249–259.

36. de Jong JW, Meijboom KE, Vanderschuren LJ, Adan RA (2013): Low control over palatable food intake in rats is associated with habitual behavior and relapse vulnerability: individual differences. PLoS One. 8:e74645.

37. Piazza PV, Deroche-Gamonet V (2013): A multistep general theory of transition to addiction. Psychopharmacology (Berl). 229:387–413.

38. Bzdok D, Meyer-Lindenberg A (2018): Machine Learning for Precision Psychiatry: Opportunities and Challenges. Biol Psychiatry Cogn Neurosci Neuroimaging. 3:223–230.

39. Pedregosa F, Varoquaux G, Gramfort A, Michel V, Thirion B, Grisel O, et al. (2011): Scikit-learn: Machine Learning in Python. J Mach Learn Res. 12:2825–2830.

40. Chollet F, others (2015): Keras. GitHub.

41. Morrow JD, Flagel SB (2016): Neuroscience of resilience and vulnerability for addiction medicine: From genes to behavior. Prog Brain Res. 223:3–18.

42. Aguilar MA, Cannella N, Ferragud A, Spanagel R (2020): Editorial: Neurobehavioural Mechanisms of Resilience and Vulnerability in Addictive Disorders. Front Behav Neurosci. 14:644495.

43. Reynolds D (2009): Gaussian Mixture Models. In: Li SZ, Jain A, editors. Encyclopedia of Biometrics. Boston, MA: Springer US, pp 659–663.

44. Forgy E (1965): Cluster analysis of multivariate data: efficiency versus interpretability of classifications. Biometrics. 21:768–769.

45. Cramer JS (2002): The Origins of Logistic Regression. Tinbergen Institute Discussion Papers.02- 119.

46. Zou J, Han Y, So SS (2008): Overview of artificial neural networks. Methods Mol Biol. 458:15–23.

47. Kumar R, Indrayan A (2011): Receiver operating characteristic (ROC) curve for medical researchers. Indian Pediatr. 48:277–287.

48. Caruana R, Lawrence S, Giles L (2000): Overfitting in neural nets: backpropagation, conjugate gradient, and early stopping. Proceedings of the 13th International Conference on Neural Information Processing Systems. Denver, CO: MIT Press, pp 381–387.

49. Puhl MD, Blum JS, Acosta-Torres S, Grigson PS (2012): Environmental enrichment protects against the acquisition of cocaine self-administration in adult male rats, but does not eliminate avoidance of a drug-associated saccharin cue. Behav Pharmacol. 23:43–53.

50. Bardo MT, Klebaur JE, Valone JM, Deaton C (2001): Environmental enrichment decreases intravenous self-administration of amphetamine in female and male rats. Psychopharmacology (Berl). 155:278–284.

51. Fung G (2001): A Comprehensive Overview of Basic Clustering Algorithms.

52. Lubke GH, Muthen B (2005): Investigating population heterogeneity with factor mixture models. Psychol Methods. 10:21–39.

53. McDermott C, Kelly JP (2008): Comparison of the behavioural pharmacology of the Lister-Hooded with 2 commonly utilised albino rat strains. Prog Neuropsychopharmacol Biol Psychiatry. 32:1816–1823.

54. Nestler EJ (2005): Is there a common molecular pathway for addiction? Nat Neurosci. 8:1445–1449.

55. Everitt BJ (1990): Sexual motivation: a neural and behavioural analysis of the mechanisms underlying appetitive and copulatory responses of male rats. Neurosci Biobehav Rev. 14:217–232.

56. Blackburn JR, Phillips AG, Jakubovic A, Fibiger HC (1989): Dopamine and preparatory behavior: II. A neurochemical analysis. Behav Neurosci. 103:15–23.

57. Murray JE, Belin D, Everitt BJ (2012): Double dissociation of the dorsomedial and dorsolateral striatal control over the acquisition and performance of cocaine seeking. Neuropsychopharmacology. 37:2456–2466.

58. Pelloux Y, Murray JE, Everitt BJ (2015): Differential vulnerability to the punishment of cocaine related behaviours: effects of locus of punishment, cocaine taking history and alternative reinforcer availability. Psychopharmacology (Berl). 232:125–134.

59. Domi A, Stopponi S, Domi E, Ciccocioppo R, Cannella N (2019): Sub-dimensions of Alcohol Use Disorder in Alcohol Preferring and Non-preferring Rats, a Comparative Study. Front Behav Neurosci. 13:3.

60. Cannella N, Cosa-Linan A, Buchler E, Falfan-Melgoza C, Weber-Fahr W, Spanagel R (2018): In vivo structural imaging in rats reveals neuroanatomical correlates of behavioral sub-dimensions of cocaine addiction. Addict Biol. 23:182–195.

61. Leggio L, Kenna GA, Fenton M, Bonenfant E, Swift RM (2009): Typologies of alcohol dependence. From Jellinek to genetics and beyond. Neuropsychol Rev. 19:115–129.

62. Kupfer DJ, First MB, Regier DA (2008): A research agenda for DSM V. American Psychiatric Pub.

63. Blanco C, Krueger RF, Hasin DS, Liu SM, Wang S, Kerridge BT, et al. (2013): Mapping common psychiatric disorders: structure and predictive validity in the national epidemiologic survey on alcohol and related conditions. JAMA Psychiatry. 70:199–208.

64. Litten RZ, Ryan ML, Falk DE, Reilly M, Fertig JB, Koob GF (2015): Heterogeneity of alcohol use disorder: understanding mechanisms to advance personalized treatment. Alcohol Clin Exp Res. 39:579–584.

65. Kwako LE, Momenan R, Litten RZ, Koob GF, Goldman D (2016): Addictions Neuroclinical Assessment: A Neuroscience-Based Framework for Addictive Disorders. Biol Psychiatry. 80:179–189.

66. Kwako LE, Schwandt ML, Ramchandani VA, Diazgranados N, Koob GF, Volkow ND, et al. (2019): Neurofunctional Domains Derived From Deep Behavioral Phenotyping in Alcohol Use Disorder. Am J Psychiatry. 176:744–753.

67. Mann K, Hermann D (2010): Individualised treatment in alcohol-dependent patients. Eur Arch Psychiatry Clin Neurosci. 260 Suppl 2:S116–120.

68. Witkiewitz K, Litten RZ, Leggio L (2019): Advances in the science and treatment of alcohol use disorder. Sci Adv. 5:eaax4043.

