## Supplementary material for "Towards a machine-learning assisted diagnosis of psychiatric disorders and their operationalization in preclinical research: evidence from studies on addiction-like behaviour in individual rats": All_alcohol_Kmean.py

import numpy as np
import pandas as pd
from sklearn.preprocessing import StandardScaler
from sklearn.cluster import KMeans

def swapping(arr):
    
    """As the labels assigned to the resilient and vulnerable rats are random, 
    we define this function to switch the labels around to always ensure that 
    1 = vulnerable and 0 = resilient. 
    
    We input the numpy array of the labels assigned by the labelencoder and the function
    returns the array with the labels swapped around if the number of 1s are greater 
    than the number 0s in the array (i.e. if the resilient rats are labelled as 1).     
    """
    
    if np.count_nonzero(arr == 1) > (len(arr) - np.count_nonzero(arr == 1)):
        
        new_arr = []
        for item in arr:
            if item == 0:
                new_arr.append(1)
            if item == 1:
                new_arr.append(0)
                
            arr = new_arr
            arr = np.asarray(arr)
    else:
        pass
    
    return(arr)

#Importing the dataset from a csv file 
dataset = pd.read_csv('All_data_alcohol.csv')
    
#Converting the rows of the pandas dataframe containing the 3 variables of interest into a numpy array
X = dataset.iloc[:,[1,2,3]].values

#Feature Scaling - INSERT A CLEARER CAPTION FOR FEATURE SCALING
sc = StandardScaler()
X = sc.fit_transform(X)

#Defining the model to contain two clusters based on our silhouette method
model = KMeans(n_clusters=2)

#Fitting the model
model.fit(X)

#Generating a list of labels from the clustering called yhat
yhat = model.predict(X)

#Applying the swap on the labels of the list of clusters if they are the wrong way round to ensure that 1 = Vulnerable
yhat = swapping(yhat)
