## Supplementary material for "Towards a machine-learning assisted diagnosis of psychiatric disorders and their operationalization in preclinical research: evidence from studies on addiction-like behaviour in individual rats": cluster_numbers_alcohol.py

# Importing the necessary libraries

from sklearn.cluster import KMeans
from sklearn.mixture import GaussianMixture
from sklearn.metrics import silhouette_score
from sklearn.preprocessing import StandardScaler
import statistics as stats
import pandas as pd


# Initializing lists to which optimal cluster number will be added for each of the 50 training and test sets
optimal_cluster_GMM_Training = []
optimal_cluster_GMM_test = []
optimal_cluster_Kmean_Training = []
optimal_cluster_Kmean_test = []


for i in range(50):
    dataset = pd.read_csv('Training'+ str(i) +'.csv') # a new training set is initialized at each loop
    X = dataset.iloc[:,[1,2,3]].values
    
    # Scaling the data to avoid skewing the weight towards the dimension with larger values
    sc = StandardScaler()
    X = sc.fit_transform(X)
    
    # defining the potential number fo clusters from which the the optimal cluster number will be chosen
    range_n_clusters = [2, 3, 4, 5, 6]
    
    # Initializing the list to which the score for each of the potential cluster number as defined
    # in 'range_n_clusters' will be added
    score = []
    
    for n_clusters in range_n_clusters:
        
        clusterer = KMeans(n_clusters=n_clusters)
        cluster_labels = clusterer.fit_predict(X)
    
        # The silhouette_score gives the average value for all the samples.
        # This gives a perspective into the density and separation of the formed
        # clusters
        silhouette_avg = silhouette_score(X, cluster_labels)
        
        score.append(silhouette_avg)
        
    # The cluster number with the maximum score is appended to the list
    optimal_cluster_Kmean_Training.append(score.index(max(score)) + 2)
    print(stats.mode(optimal_cluster_Kmean_Training)) # most common cluster number identified
    print(optimal_cluster_Kmean_Training.count(stats.mode(optimal_cluster_Kmean_Training))) # number of times the most common cluster appears
        
    
for i in range(50): 
    dataset = pd.read_csv('test'+ str(i) +'.csv') # a new test set is initialized at each loop
    X = dataset.iloc[:,[1,2,3]].values
    
    # Scaling the data to avoid skewing the weight towards the dimension with larger values
    sc = StandardScaler()
    X = sc.fit_transform(X)
    
     # defining the potential number fo clusters from which the the optimal cluster number will be chosen
    range_n_clusters = [2, 3, 4, 5, 6]
    
    # Initializing the list to which the score for each of the potential cluster number as defined
    # in 'range_n_clusters' will be added
    score = []
    
    for n_clusters in range_n_clusters:
        
        clusterer = KMeans(n_clusters=n_clusters)
        cluster_labels = clusterer.fit_predict(X)
    
        # The silhouette_score gives the average value for all the samples.
        # This gives a perspective into the density and separation of the formed
        # clusters
        silhouette_avg = silhouette_score(X, cluster_labels)
        #print(silhouette_avg)
        
        score.append(silhouette_avg)
    
    # The cluster number with the maximum score is appended to the list      
    optimal_cluster_Kmean_test.append(score.index(max(score)) + 2) # most common cluster number identified    
    print(optimal_cluster_Kmean_test.count(stats.mode(optimal_cluster_Kmean_test))) # number of times the most common cluster appears
    
for i in range(50):
    dataset = pd.read_csv('Training'+ str(i) +'.csv') # a new training set is initialized at each loop
    X = dataset.iloc[:,[1,2,3]].values
    
    # Scaling the data to avoid skewing the weight towards the dimension with larger values
    sc = StandardScaler()
    X = sc.fit_transform(X)
    
    
    range_n_clusters = [2, 3, 4, 5, 6]
    
    # Initializing the list to which the score for each of the potential cluster number as defined
    # in 'range_n_clusters' will be added
    score = []
    
    # defining the potential number fo clusters from which the the optimal cluster number will be chosen
    for n_clusters in range_n_clusters:
        
        clusterer = GaussianMixture(n_components=n_clusters)
        cluster_labels = clusterer.fit_predict(X)
    
        # The silhouette_score gives the average value for all the samples.
        # This gives a perspective into the density and separation of the formed
        # clusters
        silhouette_avg = silhouette_score(X, cluster_labels)
        #print(silhouette_avg)
        
        score.append(silhouette_avg)
    
    # The cluster number with the maximum score is appended to the list      
    optimal_cluster_GMM_Training.append(score.index(max(score)) + 2)
    print(stats.mode(optimal_cluster_GMM_Training)) # most common cluster number identified
    print(optimal_cluster_GMM_Training.count(stats.mode(optimal_cluster_GMM_Training))) # number of times the most common cluster appears
    

for i in range(50):
    dataset = pd.read_csv('test'+ str(i) +'.csv') # a new test set is initialized at each loop
    X = dataset.iloc[:,[1,2,3]].values
    
    # Scaling the data to avoid skewing the weight towards the dimension with larger values
    sc = StandardScaler()
    X = sc.fit_transform(X)
    
    # defining the potential number fo clusters from which the the optimal cluster number will be chosen
    range_n_clusters = [2, 3, 4, 5, 6]
    
    # Initializing the list to which the score for each of the potential cluster number as defined
    # in 'range_n_clusters' will be added
    score = []
    
    for n_clusters in range_n_clusters:
        
        clusterer = GaussianMixture(n_components=n_clusters)
        cluster_labels = clusterer.fit_predict(X)
    
        # The silhouette_score gives the average value for all the samples.
        # This gives a perspective into the density and separation of the formed
        # clusters
        silhouette_avg = silhouette_score(X, cluster_labels)
        #print(silhouette_avg)
        
        score.append(silhouette_avg)
     # The cluster number with the maximum score is appended to the list    
    optimal_cluster_GMM_test.append(score.index(max(score)) + 2) 
    print(stats.mode(optimal_cluster_GMM_test)) # most common cluster number identified
    print(optimal_cluster_GMM_test.count(stats.mode(optimal_cluster_GMM_test))) # number of times the most common cluster appears
