## Supplementary material for "Towards a machine-learning assisted diagnosis of psychiatric disorders and their operationalization in preclinical research: evidence from studies on addiction-like behaviour in individual rats": GMM_alcohol.py

#Importing the necessary libraries

import numpy as np
import pandas as pd
import tensorflow as tf

from sklearn.preprocessing import StandardScaler
from sklearn.model_selection import train_test_split
from sklearn.mixture import GaussianMixture
from sklearn.linear_model import LogisticRegression
from sklearn.metrics import precision_recall_fscore_support as score
from sklearn.svm import SVC
from sklearn.neighbors import KNeighborsClassifier
from sklearn.metrics import roc_auc_score

from keras import callbacks

#Defining the function 'swapping' to rightly assign 0 as resilient and 1 as vulnerable label
def swapping(arr):
    if np.count_nonzero(arr == 1) > (len(arr) - np.count_nonzero(arr == 1)):
        new_arr = []
        for item in arr:
            if item == 0:
                new_arr.append(1)
            if item == 1:
                new_arr.append(0)
                
            arr = new_arr
            arr = np.asarray(arr)
    else:
        pass
    
    return(arr)


#Initializing the lists to which the performance evaluation metrics scores are appended to.
#Accuracy, precision,recall are self-explanatory terms. AOC will have ROC AUC scores appended

accuracy_LR  = []
precision_LR = []
recall_LR    = []
AOC_LR = []

accuracy_SVM  = []
precision_SVM = []
recall_SVM    = []
AOC_SVM = []

accuracy_KNN  = []
precision_KNN = []
recall_KNN    = []
AOC_KNN = []

accuracy_ANN = []
precision_ANN = []
recall_ANN    = []
AOC_ANN = []

for i in range(50):

    #Import dataset into a pandas dataframe
    dataset = pd.read_csv('All_data_alcohol.csv')
    
    #Extracting the 3 columns of interest from the dataframe
    X = dataset.iloc[:,[1,2,3]].values
    
    # Splitting the dataset into the Training set and Test set.
    #The random_state = i will ensure that the current value of i will be used
    #to do the train-test split. This will happen 50 times with
    #the i value ranging from 0 to 49
    X_train, X_test = train_test_split(X, test_size = 0.33, random_state = i) 
    
    # Feature Scaling (it is done to avoid large influences of one variable on the model if it has larger values)
    sc = StandardScaler()
    X_train = sc.fit_transform(X_train)
    X_test = sc.transform(X_test)
    
        
    #UNSUPERVISED LEARNING METHOD - Clustering the training dataset using the Gaussian Mixture Model 
    
    # define the model
    model = GaussianMixture(n_components=2)
    # fit the model
    model.fit(X_train)
    # assign a cluster to each example
    yhat = model.predict(X_train)
    yhat = swapping(yhat)
    
    
    #UNSUPERVISED LEARNING METHOD - Clustering the test dataset using the Gaussian Mixture Model 
    
    # define the model
    model = GaussianMixture(n_components=2)
    # fit the model
    model.fit(X_test)
    # assign a cluster to each example
    yhattest = model.predict(X_test)
    yhattest = swapping(yhattest)
    
    
    
    #SUPERVISED LEARNING METHOD 1 - Training the LOGISTIC REGRESSION model on the Training set
    classifierLR = LogisticRegression(random_state = 0)
    classifierLR.fit(X_train, yhat)
    
    #Predicting the labels for the X_test raw data inputs
    y_predLR = classifierLR.predict(X_test)
    y_predLR = swapping(y_predLR)
    
    #Calculating the performance evaluation metrics for Logistic Regression  
    precision,recall,fscore,support=score(yhattest, y_predLR,average='macro')
    accuracy_LR.append(fscore)
    precision_LR.append(precision)
    recall_LR.append(recall)
    AOC_LR.append(roc_auc_score(yhattest, y_predLR))
    
    
    
    #SUPERVISED LEARNING METHOD 2 - Training the Support Vector Machine Classifier with rbf kernel on the TRaining set
    classifier_SVM = SVC(kernel = 'rbf', random_state = 0)
    classifier_SVM.fit(X_train, yhat)
    
    #Predicting the labels for the X_test raw data inputs
    y_predSVM = classifier_SVM.predict(X_test)
    y_predSVM = swapping(y_predSVM)
    
    #Calculating the performance evaluation metrics for Support Vector Machine
    precision,recall,fscore,support=score(yhattest, y_predSVM,average='macro')
    accuracy_SVM.append(fscore)
    precision_SVM.append(precision)
    recall_SVM.append(recall)
    AOC_SVM.append(roc_auc_score(yhattest, y_predSVM))
    
    
    
    #SUPERVISED LEARNING METHOD 3 - Training the K-NN model on the Training set
    classifierKNN = KNeighborsClassifier(n_neighbors = 5, metric = 'minkowski', p = 2)
    classifierKNN.fit(X_train, yhat)
    
    #Predicting the labels for the X_test raw data inputs
    y_predKNN = classifierKNN.predict(X_test)
    y_predKNN = swapping(y_predKNN) 
    
    #Calculating the performance evaluation metrics for k-Nearest Neighbours
    precision,recall,fscore,support=score(yhattest, y_predKNN,average='macro')
    accuracy_KNN.append(fscore)
    precision_KNN.append(precision)
    recall_KNN.append(recall)
    AOC_KNN.append(roc_auc_score(yhattest, y_predKNN))
    
    
    
    #SUPERVISED LEARNING METHOD 4 - Training Artificial neural networks
    N, D = X_train.shape
      
    model = tf.keras.models.Sequential([
    tf.keras.layers.Dense(500, activation= 'elu'),# the number of hidden layers tested are 5, 50, 500
    tf.keras.layers.Dense(units= 6),
    tf.keras.layers.Dropout(0.2),
    tf.keras.layers.Dense(2, input_shape = (D,), activation='softmax')
    ])
      
    #Compile the model
    model.compile(optimizer = 'adam', loss = 'sparse_categorical_crossentropy', metrics = ['accuracy'])
    earlystopping = callbacks.EarlyStopping(monitor ="val_accuracy", 
                                        mode ="max", patience = 25, 
                                        restore_best_weights = True)
    
    r = model.fit(X_train, yhat, validation_data= (X_test, yhattest), epochs=100, callbacks = [earlystopping])
    
    #Calculating the performace evaluation metrics for Artificial neural networks
    a = model.evaluate(X_test, yhattest)[1]
    accuracy_ANN.append(a)
    
    P = model.predict(X_test)
    P = np.round(P[:,1]).flatten()
    
    m = tf.keras.metrics.Recall()
    m.update_state(P, yhattest)
    recall_ANN.append(m.result().numpy())
    
    n = tf.keras.metrics.Precision()
    n.update_state(P, yhattest)
    precision_ANN.append(n.result().numpy())
    
    AOC_ANN.append(roc_auc_score(yhattest, P))
