## Supplementary material for "Towards a machine-learning assisted diagnosis of psychiatric disorders and their operationalization in preclinical research: evidence from studies on addiction-like behaviour in individual rats": SOM

### Supplementary Methods:

The present analysis uses total responses performed by the rats while in the original experiments for addiction-like behavior for cocaine, compulsivity was expressed as a percentage of baseline, while the last completed ratio in the progressive-ratio test was considered to assess motivation.

GMM clustering (1) was used to mimic the bimodal threshold logic used in the 3-Criteria model since it is ideal for determining clusters when the central limit theorem cannot be applied and the data distribution shows bi/multimodal distribution. K-mean/K-median algorithms were used for their relative ease of implementation, their ability to identify clusters of different shapes and sizes as well as their scalability (2).

K nearest neighbor generates a mathematical model for the rats of the TRAINING SET behavioral data and the labels assigned by the clustering algorithm based on the KNN rule that states that similar data points are closer and hence belong together (2). By submitting the behavioral data of the TEST set rats to this mathematical model, their labels are predicted based on how closely they resemble the TRAINING SET data points.

The logit function used in Logistic regression generates a mathematical model by mapping between 0 and 1 the behavioral data of the TRAINING SET rats and the labels assigned by the clustering algorithm. When the behavioral data of the TEST SET rats is submitted to this mathematical model, their labels as Resilient or Vulnerable are predicted by where they fall on the linear equation output between 0 and 1 with the cut off being 0.5 (3).

Support Vector Machines generate a mathematical model from the behavioral data of TRAINING SET rats and the labels assigned by clustering algorithms by identifying a hyperplane in a multidimensional vector space such that the distance between the nearest points of the two groups from the hyperplane is maximized. When the behavioral data of the TEST SET rats is submitted to this mathematical model, the vulnerability label of each rat is determined by their location in the vector space with respect to the hyperplane (4).

Finally, supervised Artificial Neural Networks take the behavioral data of the TRAINING SET rats and pass it through a feed-forward network of hidden neurons starting with random weights to calculate a 'cost function' eventually to assign labels as resilient and vulnerable to these rats. Back-propagation adjusts these weights to minimize this 'cost function' and adjust these labels to get as close to the labels of the TRAINING SET rats as assigned by the clustering algorithm as possible thereby generating a mathematical model (**Figure 3**).

Accuracy: The proportion of rats from the TEST SET correctly predicted by the Supervised algorithms compared to the labels given by the clustering algorithms

**TV + TR**

**TV + TR + FV + FR**

**Accuracy =**

Precision: Of all the rats predicted as Vulnerable by the Supervised algorithms, how many were also labelled as Vulnerable by the Clustering algorithms. The higher the precision, lower is the proportion of rats falsely predicted as vulnerable

**TV**

**TV + FV**

**Precision =**

Recall (Sensitivity): Of all the rats labelled as Vulnerable by the Clustering algorithm, how many were correctly identified as Vulnerable by the Supervised Classification algorithm

**TV**

**TV + FR**

**Recall =**

AUC ROC score: Receiver Operating Characteristics (ROC) curve (5) plots the False Vulnerable rate [FV/(TR+FV)](1-precision) on the X-axis versus the True Vulnerable rate [TV/(TV+FR)] on the y-axis for several candidate threshold values between 0 and 1. The AUC ROC score is a useful tool to compare different classification models since it demonstrates the change in the relationship between precision and recall by varying the threshold to identify a TV rat.

### References:

1.	Reynolds D (2009): Gaussian Mixture Models. In: Li SZ, Jain A, editors. *Encyclopedia of Biometrics*. Boston, MA: Springer US, pp 659-663.

2.	Forgy E (1965): Cluster analysis of multivariate data : efficiency versus interpretability of classifications. *Biometrics*. 21:768-769.

3.	Cramer JS (2002): The Origins of Logistic Regression. *Tinbergen Institute Discussion Papers*.02-119.

4.	Pedregosa F, Varoquaux G, Gramfort A, Michel V, Thirion B, Grisel O, et al. (2011): Scikit-learn: Machine Learning in Python. *J Mach Learn Res*. 12:2825–2830.

5.	Kumar R, Indrayan A (2011): Receiver operating characteristic (ROC) curve for medical researchers. *Indian Pediatr*. 48:277-287.
